# The role of diffusion and perivascular spaces in dynamic susceptibility contrast MRI

**DOI:** 10.1101/614008

**Authors:** Jonathan Doucette, Christian Kames, Enedino Hernández-Torres, Anthony Traboulsee, Alexander Rauscher

**Author notes:** **Corresponding Author** Jonathan Doucette, PhD Student, Department of Physics and Astronomy, UBC MRI Research Centre, University of British Columbia, M10 - Purdy Pavilion, 2221 Wesbrook Mall, Vancouver, BC V6T 2B5. Parts of this work have been presented as an oral presentation at ISMRM 2018.

## Abstract

We investigated the effects of brain tissue orientation, diffusion, and perivascular spaces on dynamic susceptibility contrast MRI. A 3D numerical model of a white matter voxel was created that consists of an isotropic capillary bed and anisotropic vessels that run in parallel with white matter tracts and are surrounded by perivascular spaces. The signal within the voxel was simulated by solving the Bloch-Torrey equation. Experimental perfusion data were acquired with a gradient echo dynamic susceptibility contrast scan. White matter fibre orientation was mapped with diffusion tensor imaging. Our numerical model of the contrast agent induced increase in R_2_^*^, as a function of tissue orientation, was fit to dynamic susceptibility contrast MRI data from thirteen subjects by minimizing the bias-corrected Akaike information criterion. White matter blood volume fraction in both the isotropic and the anisotropic vessels was determined as a free parameter, and results were analyzed as a function of diffusivity and perivascular space size. Total white matter blood volume was found to be 2.57%, with one third of the blood residing in blood vessels that run parallel with white matter tracts. Gradient echo dynamic susceptibility contrast MRI strongly depended on white matter tissue orientation and, according to the numerical simulations, this effect is amplified by diffusion and perivascular spaces.

## Introduction

In dynamic susceptibility contrast (DSC) MRI, a paramagnetic contrast agent (CA) is injected during the acquisition of a rapid T_2_^*^ or T_2_ weighted scan. The modification of the signal due to the presence of the CA within the vascular system allows for the computation of cerebral blood flow (CBF) and volume (CBV). In the brain’s white matter (WM), the vascular architecture consists of a structurally isotropic network of small vessels and larger vessels that, on average, run in parallel with the WM tracts^1–3^. The resulting vascular anisotropy causes gradient-echo (GRE) DSC MRI and the maps of cerebral blood flow and volume derived from this scan to exhibit a strong orientation dependency^4^. This phenomenon is caused by the orientation dependency of the field inhomogeneities that are created by the blood vessels and amplified by the paramagnetic CA. The field inhomogeneities around a vessel become stronger with increasing angle α between the vessel and B_0_. Therefore, the change in R_2_^*^created by a contrast agent not only depends on the CA concentration but also on tissue orientation. The orientation effect on the CA induced change in R_2_^*^ (ΔR_2_^*^), as well as on CBF and CBV derived from ΔR_2_^*^, is on the order of 100%. Modeling the vascular architecture and signal loss due to static dephasing within the field inhomogeneities created by the vessels showed that the average blood volume in white matter is 2%, and that half of the blood resides in the anisotropic component of the vascular tree. While a model based on static dephasing fits the measured data reasonably well, diffusion of spins within the magnetic field inhomogeneities introducing further signal loss should be taken into account^5^. Furthermore, larger vessels are surrounded by perivascular spaces (PVS), which are filled with fluid and therefore have high diffusion coefficients and long T_2_ relaxation times^6^, which means that these PVS influence the DSC signal. Here, we investigate the extent to which diffusion and PVS impact the measured ΔR_2_^*^ value as a function of WM fibre orientation in GRE DSC. We explore how the inclusion of diffusion and PVS into the fibre orientation dependent GRE DSC model influences the determined vascular parameters and the goodness of fit. In addition, we extend the 2D simulation used in a previous study of GRE DSC^4^ to the simulation of a 3D voxel in order to fully capture the effects of diffusion within the complex 3D anisotropic vascular architecture.

## Materials and Methods

### Data acquisition

This study was approved by the Clinical Research Ethics Board of our institution (H12-01153, 20 June 2012) and is in accord with the Declaration of Helsinki. All subjects gave written informed consent. Subjects and data acquisition are explained in detail in a previous publication for the same cohort^4^. In short, DSC (TR/TE = 2417/40 ms; 40 dynamics, reconstructed voxel size = 1.75 × 1.75 × 4 mm^3^; Magnevist 0.2 ml/kg body weight, 5 ml/s followed by 20 ml saline flush) and diffusion tensor imaging (DTI) data (TR/TE = 5640/75 ms; b = 1000, 32 directions, reconstructed voxel size = 1.88 × 1.88 × 2.5) from thirteen subjects with multiple sclerosis (MS) were acquired on a 3T scanner (Philips Achieva) equipped with an 8-channel SENSE head coil. Perfusion maps were registered to DTI using the linear image registration tool FLIRT from the FSL software package^7^. A WM mask was created using FSL’s FAST tool on the combination T_1_T_2_ = (T_1_ − T_2_)/ (T_1_ + T_**2**_) of a 3D T_1_-weighted sequence (TR/TE = 7.6/3.7 ms; reconstructed voxel size = 0.8 × 0.8 × 0.8 mm^3^) and a 3D T_2_-weighted sequence (TR/TE = 2500/363 ms; reconstructed voxel size = 0.8 × 0.8 × 0.8 mm^3^). Maps of local fibre orientation were computed from the DTI data as described previously^8^. Following motion correction of the DSC data, the change in R_2_^*^ due to the contrast agent was calculated according to

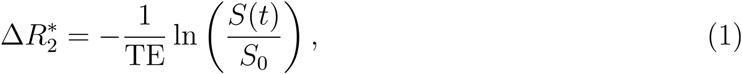

where TE is the echo time, S(t) is the signal at any time during the DSC experiment, for example at peak contrast agent concentration, and S_0_ is the signal without contrast agent^9,10^. The ΔR_2_^*^ value for the peak contrast agent concentration was determined by taking the value at the time point with the lowest signal for each voxel, i.e. by computing a minimum intensity projection along the time axis of the motion corrected data. Then, ΔR_2_^*^ in each WM voxel was sorted according to fibre angle α into bins of width 5^∘^ centred on angles ranging from 2.5^∘^ to 87.5^∘^.

### Numerical model and parameter fit

Simulation of the transverse magnetization was performed within a 3D voxel containing isotropic and anisotropic blood vessels. The mean radius of the small isotropically oriented vessels was 7 μm^11^. The large vessels were variable in number and size and ran in parallel with the WM direction. The total blood volume fraction BVF determines the total number of vessels present, with an amount iBVF contained in the isotropic vasculature, and the remaining aBVF = BVF – iBVF contained in the anisotropic vasculature. Perivascular spaces were included which surround the large anisotropic vessels. These spaces are modeled as a hollow cylinders with inner radii equal to the anisotropic vessel radii and variable outer radii; the outer radius relative to the inner radius is denoted ρ, and the volume of the PVS relative to the contained anisotropic vessel is then given by ν = ρ^2^−1. Field inhomogeneities for a given vascular configuration were calculated by convolving the magnetic susceptibility distribution associated with the vascular tree with a unit dipole^12^. For the magnetic susceptibility of the contrast agent a value of 0.34 ppm/mM was used^13^. Within the voxel geometry, the transverse magnetization is propagated forward in time by solving the Bloch-Torrey equation

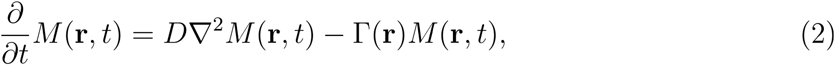

up until the DSC scan’s echo time of 40 ms, as described in detail in a previous study^14^. Here, M is the complex transverse magnetization, D is the diffusion coefficient or diffusivity, and Γ(**r**) is the complex decay rate given by R_2_(**r**) + iω(**r**), with ω(**r**) being the local resonance frequency determined by the magnetic field. The parameters iBVF, aBVF, and the peak contrast agent concentration CA were optimized using weighted non-linear regression with MATLAB’s *fmincon* function, where the model residuals were weighted by the fraction of data points contained in each WM fibre orientation interval. The loss function applied to the weighted residuals which was minimized was the bias-corrected Akaike information criterion (AICc), as it performs better as a measure of comparing nonlinear models than traditional metrics such as the coefficient of determination (R^2^)^15^. The minimization was performed for a variable number of large vessels ranging from N = 1 to N = 9 with a diffusivity of D = 2 μm^2^ms^−1^ and a PVS volume of ν = 1X. To test the effects of changes in diffusion constant, simulations were performed for diffusivities ranging from D = 0 μm^2^ms^−1^ to D = 3 μm^2^ms^−1^ using the vascular geometry determined from the D = 2 μm^2^ms^−1^ and ν = 1X minimization. Similarly, simulations were performed with the same geometry, D = 2 μm^2^ms^−1^, and ν varying from 0X to 10X. The simulation results are shown in Figures 1B-E, with an example geometry voxel shown in Figure 1A.

## Results

For static dephasing only and without PVS, i.e. D = 0 μm^2^ms^−1^ and ν = 0X, the best fit resulted in N = 1 anisotropic vessels, iBVF = 1.62%, aBVF = 0.84%, and a peak contrast agent concentration of 3.90 mM. Inclusion of diffusion with D = 2 μm^2^ms^−1^ and PVS with relative volume ν = 1X improved the model fit with the best fit resulting in N = 4 anisotropic vessels with corresponding radii of 43.7 μm, iBVF = 1.72%, aBVF = 0.85%, and a peak contrast agent concentration of 3.65 mM. The goodness of fit improved, as demonstrated by a decrease in AICc from −36.47 to −66.91; for comparison, R^2^ increased from 0.951 to 0.991. Holding PVS volume constant at ν = 1X and varying the diffusivity from D = 0 μm^2^ms^−1^ to D = 3 μm^2^ms^−1^ showed increasing ΔR values for all angles α, with the largest changes of 0.7 Hz occurring near α = 0° and α = 90° and the smallest changes of 0.3 Hz occurring near α = 45°. Holding diffusivity constant at D = 2 μm^2^ms^−1^ and varying the PVS volume from ν = 0X to ν = 10X showed ΔR_2_^*^ increases which increase with α. The effect of PVS is modest for small volumes below ν = 3X, but quickly becomes large for larger volumes, with a maximal increase of 1.5 Hz occurring at α = 90° with ν = 10X. Detailed tables of all simulation results, including fit parameters and goodness of fit measures, are presented in the Supplementary Figure 1.

**Figure 1.**
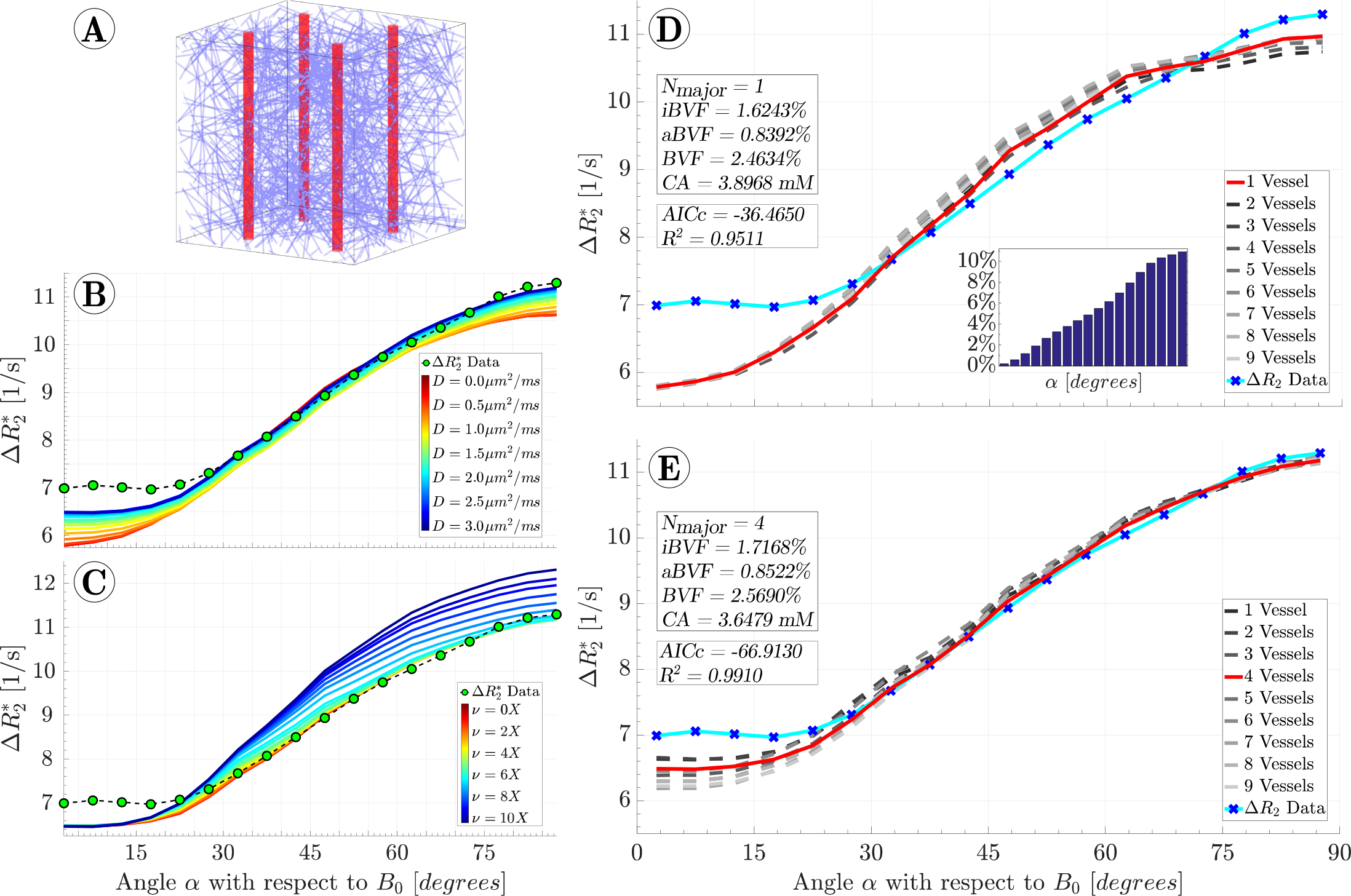
A) Example of a simulated voxel populated with an isotropic vascular bed and 4 anisotropic vessels. B) Simulations with the diffusivity varying from D = 0 μm^2^ms^−1^ to D = 3 μm^2^ms^−1^, showing that increases in D lead to increases in ΔR_2_^*^. The vascular parameters resulting from the fit in Figure 1D were used for each simulation. C) Simulations with the PVS volume varying from ν = 0X to ν = 10X, using the same vascular parameters as in (B) and a constant D of 2 μm^2^ms^−1^, showing that increased PVS volume also leads to increased ΔR_2_^*^. D) The best fit obtained with static dephasing. The histogram insert shows the percentage of voxels contained in each orientation interval which were used as weights in all fitting procedures. Note that the distribution of the number of voxels in each interval is due to a geometric effect: the fraction of uniformly randomly oriented vectors with angle α with respect to B_0_ is proportional to sin(α). E) The best fit obtained when diffusion is included in the model results in a total blood volume of 2.57%. About one third of the blood resides in vessels that run in parallel with white matter tracts.

## Discussion

We extended previously published work for gradient echo DSC by incorporating the effects of diffusion and perivascular spaces, as well as by simulating a 3D voxel instead of a 2D slice of a voxel. Diffusion was found to increase the simulated ΔR_2_^*^value independently of the orientation of the WM fibres with respect to B_0_, providing further confirmation that diffusion mediated signal loss is dominated by isotropic vasculature as opposed to anisotropic vasculature. However, the effect of diffusion around the larger anisotropic vessels is further augmented by the fluid within PVS, which has a high diffusion coefficient. This finding agrees well with the optimized parameter values of iBVF = 1.72% and aBVF = 0.85%, indicating that approximately two-thirds of WM vasculature is comprised by isotropic vessels. The total blood volume fraction in WM associated with the best fit was 2.57%, agreeing well with literature values, which report a white matter blood volume ranging from 1.3% to 2.6%^16–18^, and also with the 2D simulation of static dephasing, where a total blood volume of 2% was reported^4^. In particular, our result is in excellent agreement with the total blood volume of 2.6% determined with positron emission tomography^16^, which is considered the gold standard for cerebral blood volume measurements. With neither diffusion nor PVS included in the model, the total blood volume was found to be 2.46%. Inclusion of diffusion and PVS improved the goodness of fit of the model, as measured by both AICc and R^2^. Nevertheless, the vascular parameters computed with the extended model are not very different from the static dephasing model, since the increase in R_2_^*^ due to the contrast agent is dominated by static dephasing. However, a static dephasing model is not able to estimate the calibre of the anisotropic vessels. With diffusion, the size of these vessels starts to play a role, because diffusion has a larger effect around small vessels than around large vessels. Surprisingly little is known about the architecture and function of PVS. For example, very recent work demonstrated that PVS shrink by 90% during fixation^19^. We are aware of only one study where the effects of PVS on quantitative MRI were investigated. Sepehrband et al. showed that diffusion tensor imaging measures, such as fractional anisotropy and mean diffusivity, are biased by PVS^20^. These authors demonstrate that this effect is due to the high diffusivity of the fluid within the PVS, and to the restricted diffusion of PVS fluid in the direction perpendicular to the vessel axis.

It should be emphasized that for non-linear models, R^2^ is not the best measure of goodness of fit. Nevertheless, it is often reported even for non-linear models. Here, we report R^2^ in addition to AICc because it allows for a more intuitive comparison of fit quality. In the previous work in the same cohort, using 2D simulation and static dephasing, we obtained a lower total blood volume of 2%, but peak contrast agent concentration was higher at 7.6 mM and the anisotropic vascular volume was 50% of the total volume ^4^. Using spin echo DSC data obtained in a different cohort of healthy volunteers, we found that 56% of the blood is contained in the anisotropic component of the vascular tree^14^, similar to the 66% found in this work. It should be noted that in the previous study on static dephasing we used a radius of 13.7 μm for the isotropic vasculature, which was obtained from a study on vessel size imaging with MRI^21^. However, we believe that the value of 7 μm determined directly with microscopy is more reliable^11^.

The data collected for this study was from patients with multiple sclerosis. Due to the administration of contrast agent, this study could not have been performed on healthy patients. Compared to perfusion measurements arising from e.g. stroke or tumour, MS has the smallest impact on the vascular architecture. Nevertheless, since MS is a neurodegenerative disease that alters myelin, one may naturally be concerned with inferring WM fibre tract orientation from DTI of MS patients. However, this issue is a minor one for several reasons. First, regions containing MS lesions comprise only a small fraction of the total WM volume. Even the orientation dependency of the R_2_^*^ decay, which is directly influenced by myelin, does depend on whether MS lesions are included in the analysis or not^8^. Second, myelination of WM tracts is not required for DTI in the first place. For example, Figure 1 in a study by Vinall et al.^22^ shows a map of principal diffusion direction weighted with the fractional anisotropy acquired in an infant at 30 weeks postmenstrual age, i.e. approximately 10 weeks premature. All prominent WM structures are clearly visible and their orientations are well defined, despite little to no myelination of the WM tracts at this age.

Multiple sclerosis may also increase the relative size of the PVS due WM atrophy ^23^, and PVS size is also known to increase with age^24^. Moreover, CBF and CBV were generally found to be reduced in the normal appearing white matter in MS^25^. While the finding that PVS have an influence on DSC measurements remains valid, the size of the PVS estimated herein should therefore not be generalized to other populations. Due to the orientation dependence of DSC, previous findings in WM in patient studies should be interpreted with caution. In particular CBF and CBV in MS lesions, which are usually traversed by a vein^26^, may be strongly influenced by the vein’s orientation.

One limitation of the current model is that water molecules are allowed to freely move across the boundary between outside and inside the vessels. This simplification may explain the large differences in ΔR_2_^*^ between static dephasing and the extended model at small angles, as this is where the difference in field strength between interior and exterior of the vessels is largest. Note, however, that the weights at low angles are small due to the small number of voxels contributing to these data. Therefore, fits are relatively insensitive to the behaviour at small angles. Moreover, since vessels occupy less than 3% of the tissue volume, the error caused by this simplification is small. It should also be kept in mind that this approach only provides information on average WM vasculature. There will be voxels with considerably more or less vascular anisotropy than this average. Furthermore, voxels which have low fractional anisotropy and/or contain crossing fibres have a less well defined local fibre orientation. However, since WM voxels were pooled from across the entire WM and averaged, such WM tract specific differences in vascular architecture are averaged away.

In summary, we show that the effects of the paramagnetic contrast agent on the MR signal are modulated by the anisotropic vascular architecture and by diffusion within perivascular spaces, resulting in a strong orientation dependency of gradient echo DSC.

## Author Contributions

JD: data analysis, numerical simulations, interpretation of results, writing of manuscript; CK: numerical simulations; EHT: data analysis; AT: data acquisition, interpretation of results; AR: idea, study supervision, interpretation of results, writing of manuscript.

## Acknowledgments and Funding

This work was funded by the Natural Sciences and Engineering Research Council of Canada (016-05371) and the National Multiple Sclerosis Society (RG-1507-05301). JD was funded by the Natural Sciences and Engineering Research Council of Canada (USRA-497681-2016). AR is funded by Canada Research Chairs. All raw data and code are available from the authors upon request.

## Abbreviations

(WM): White Matter
(PVS): Perivascular Space
(DSC): Dynamic Susceptibility Contrast
(CA): Contrast Agent
(aBVF): Anisotropic Blood Volume Fraction
(aBVF): Isotropic Blood Volume Fraction

---

**Supplemental Figure 1.**
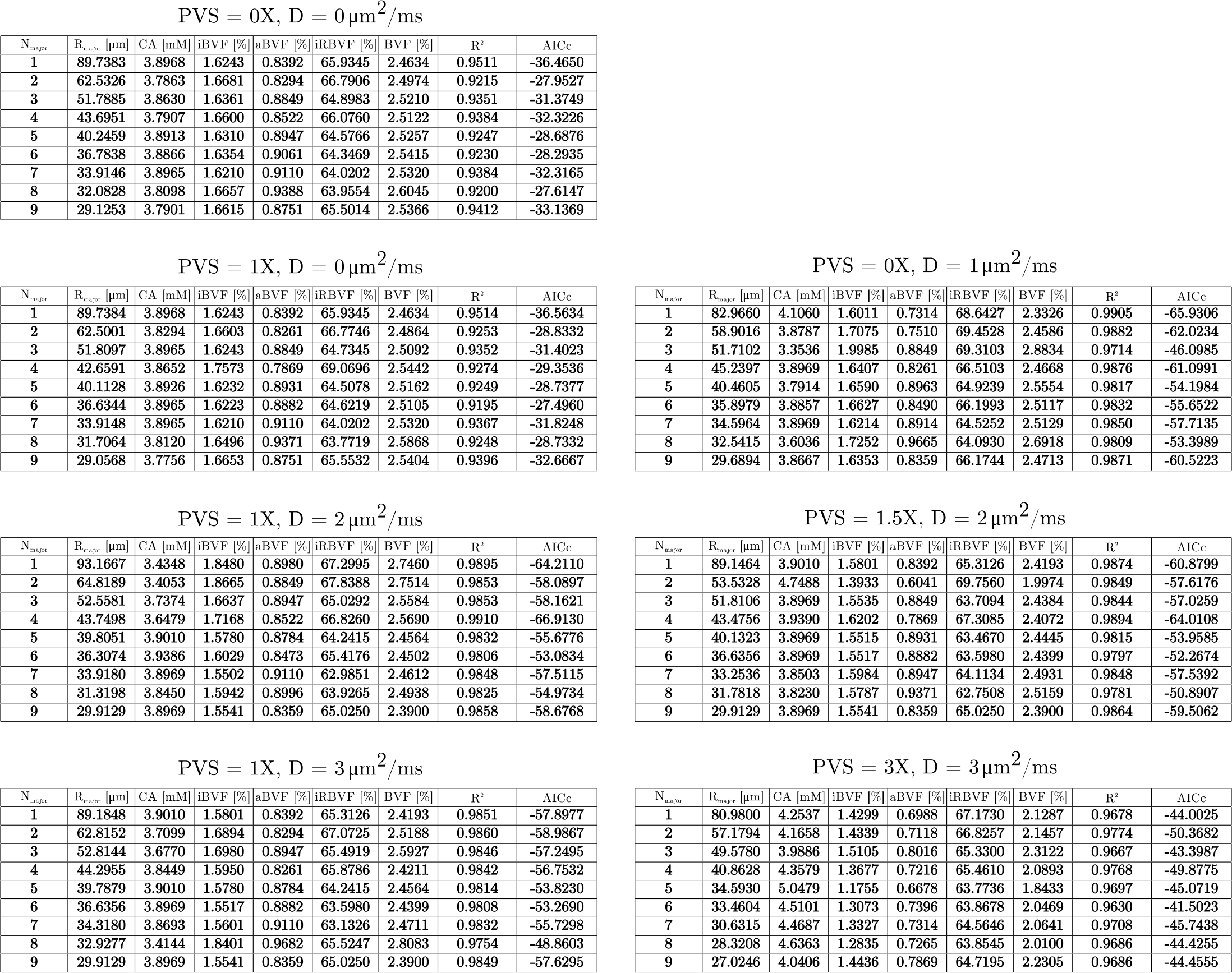
Detailed tables of minimization experiments. Diffusion coefficients and perivascular space volumes were varied between tables. Within tables, minimization results are shown for the number of anisotropic vessels varying from N = 1 to N = 9. In order from left to right, the columns are labels represent: the number of anisotropic vessels, N_major_; the radius of the anisotropic vessels, R_major_; the contrast agent concentration, CA; the isotropic blood volume fraction, iBVF; the anisotropic blood volume fraction, aBVF; the relative isotropic blood volume fraction, iRBVF = iBVF/BVF; the total blood volume fraction, BVF = iBVF + aBVF; the coefficient of determination of the resulting fit, R^2^; the bias-corrected Akaike information criterion of the resulting fit, AICc.

## References

1. Nonaka H, Akima M, Hatori T, Nagayama T, Zhang Z, Ihara F. Microvasculature of the human cerebral white matter: Arteries of the deep white matter. Neuropathology. 2003;23(2):111–118. doi:10.1046/j.1440-1789.2003.00486.x

2. Nonaka H, Akima M, Hatori T, Nagayama T, Zhang Z, Ihara F. The microvasculature of the cerebral white matter: arteries of the subcortical white matter. J Neuropathol Exp Neurol. 2003;62(2):154–161.

3. Okudera T, Huang YP, Fukusumi A, Nakamura Y, Hatazawa J, Uemura K. Micro-angiographical studies of the medullary venous system of the cerebral hemisphere. Neuropathology. 1999;19(1):93–111. doi:10.1046/j.1440-1789.1999.00215.x

4. Hernández-Torres E, Kassner N, Forkert ND, et al. Anisotropic cerebral vascular architecture causes orientation dependency in cerebral blood flow and volume measured with dynamic susceptibility contrast magnetic resonance imaging. J Cereb Blood Flow Metab. June 2016:0271678X16653134. doi:10.1177/0271678X16653134

5. Boxerman JL, Hamberg LM, Rosen BR, Weisskoff RM. Mr contrast due to intravascular magnetic susceptibility perturbations. Magn Reson Med. 1995;34(4):555–566. doi:10.1002/mrm.1910340412

6. Kwee RM, Kwee TC. Virchow-Robin Spaces at MR Imaging. RadioGraphics. 2007;27(4):1071–1086. doi:10.1148/rg.274065722

7. Jenkinson M, Beckmann CF, Behrens TEJ, Woolrich MW, Smith SM. FSL. NeuroImage. 2012;62(2):782–790. doi:10.1016/j.neuroimage.2011.09.015

8. Hernández-Torres E, Wiggermann V, Hametner S, et al. Orientation Dependent MR Signal Decay Differentiates between People with MS, Their Asymptomatic Siblings and Unrelated Healthy Controls. PLOS ONE. 2015;10(10):e0140956. doi:10.1371/journal.pone.0140956

9. Østergaard L, Weisskoff RM, Chesler DA, Gyldensted C, Rosen BR. High resolution measurement of cerebral blood flow using intravascular tracer bolus passages. Part I: Mathematical approach and statistical analysis. Magn Reson Med. 1996;36(5):715–725. doi:10.1002/mrm.1910360510

10. Østergaard L, Chesler DA, Weisskoff RM, Sorensen AG, Rosen BR. Modeling Cerebral Blood Flow and Flow Heterogeneity from Magnetic Resonance Residue Data. J Cereb Blood Flow Metab. 1999;19(6):690–699. doi:10.1097/00004647-199906000-00013

11. Meier-Ruge W, Hunziker O, Schulz U, Tobler H-J, Schweizer A. Stereological changes in the capillary network and nerve cells of the aging human brain. Mech Ageing Dev. 1980;14(1):233–243. doi:10.1016/0047-6374(80)90123-2

12. Marques JP, Bowtell RW. Using forward calculations of the magnetic field perturbation due to a realistic vascular model to explore the BOLD effect. NMR Biomed. 2008;21(6):553–565. doi:10.1002/nbm.1224

13. Weisskoff RM, Kiihne S. MRI susceptometry: Image-based measurement of absolute susceptibility of MR contrast agents and human blood. Magn Reson Med. 1992;24(2):375–383. doi:10.1002/mrm.1910240219

14. Doucette J, Wei L, Hernández-Torres E, et al. Rapid solution of the Bloch-Torrey equation in anisotropic tissue: Application to dynamic susceptibility contrast MRI of cerebral white matter. NeuroImage. 2019;185:198–207. doi:10.1016/j.neuroimage.2018.10.035

15. Spiess A-N, Neumeyer N. An evaluation of R2 as an inadequate measure for nonlinear models in pharmacological and biochemical research: a Monte Carlo approach. BMC Pharmacol. 2010;10:6. doi:10.1186/1471-2210-10-6

16. Leenders KL, Perani D, Lammertsma AA, et al. Cerebral blood flow, blood volume and oxygen utilization. Normal values and effect of age. Brain J Neurol. 1990;113 (Pt 1):27–47.

17. Helenius J, Perkiö J, Soinne L, et al. Cerebral Hemodynamics in a Healthy Population Measured by Dynamic Susceptibility Contrast Mr Imaging. Acta Radiol. 2003;44(5):538–546. doi:10.1080/j.1600-0455.2003.00104.x

18. Arakawa S, Wright PM, Koga M, et al. Ischemic Thresholds for Gray and White Matter. Stroke. 2006;37(5):1211–1216. doi:10.1161/01.STR.0000217258.63925.6b

19. Mestre H, Tithof J, Du T, et al. Flow of cerebrospinal fluid is driven by arterial pulsations and is reduced in hypertension. Nat Commun. 2018;9(1):4878. doi:10.1038/s41467-018-07318-3

20. Sepehrband F, Cabeen RP, Choupan J, et al. Perivascular space fluid contributes to diffusion tensor imaging changes in white matter. bioRxiv. January 2019:395012. doi:10.1101/395012

21. Jochimsen TH, Ivanov D, Ott DVM, et al. Whole-brain mapping of venous vessel size in humans using the hypercapnia-induced BOLD effect. NeuroImage. 2010;51(2):765–774. doi:10.1016/j.neuroimage.2010.02.037

22. Vinall J, Grunau RE, Brant R, et al. Slower Postnatal Growth Is Associated with Delayed Cerebral Cortical Maturation in Preterm Newborns. Sci Transl Med. 2013;5(168):168ra8–168ra8. doi:10.1126/scitranslmed.3004666

23. Sastre-Garriga J, Pareto D, Rovira À. Brain Atrophy in Multiple Sclerosis: Clinical Relevance and Technical Aspects. Neuroimaging Clin N Am. 2017;27(2):289–300. doi:10.1016/j.nic.2017.01.002

24. Laveskog A, Wang R, Bronge L, Wahlund L-O, Qiu C. Perivascular Spaces in Old Age: Assessment, Distribution, and Correlation with White Matter Hyperintensities. AJNR Am J Neuroradiol. 2018;39(1):70–76. doi:10.3174/ajnr.A5455

25. Lapointe E, Li DKB, Traboulsee AL, Rauscher A. What Have We Learned from Perfusion MRI in Multiple Sclerosis? AJNR Am J Neuroradiol. 2018;39(6):994–1000. doi:10.3174/ajnr.A5504

26. Sati P, Oh J, Constable RT, et al. The central vein sign and its clinical evaluation for the diagnosis of multiple sclerosis: a consensus statement from the North American Imaging in Multiple Sclerosis Cooperative. Nat Rev Neurol. 2016;12(12):714–722. doi:10.1038/nrneurol.2016.166

